# The Gene Encoding Ornithine Decarboxylase for Putrescine Biosynthesis Is Essential for the Viability of *Fusobacterium nucleatum*

**DOI:** 10.1101/2025.09.02.673652

**Authors:** Shiqi Xu, Bibek G C, Alex Phan, Chenggang Wu

## Abstract

*Fusobacterium nucleatum* is a Gram-negative anaerobe associated with periodontitis and colorectal cancer. It secretes putrescine, a polyamine that promotes biofilm formation by oral co-colonizers and enhances the proliferation of cancer cells. However, the physiological importance of putrescine for *F. nucleatum* itself remains unexplored. Here, we show that putrescine biosynthesis, mediated by the ornithine decarboxylase gene *oda*, is essential for *F. nucleatum* viability. Deletion of *oda* was only possible when a functional copy was provided in trans, and CRISPR interference of *oda* expression resulted in complete growth arrest. The essentiality of *oda* was conserved across multiple subspecies. Supplementation with exogenous putrescine enabled the isolation of a conditional *oda* mutant whose growth was strictly putrescine-dependent. Putrescine depletion caused filamentation, membrane disruption, detergent hypersensitivity, and lysis in hypoosmotic conditions, indicating a critical role in maintaining cell envelope integrity. RNA sequencing revealed broad transcriptional remodeling under putrescine-limited conditions, including upregulation of genes involved in lipid metabolism, osmoprotection, and cell wall remodeling. Notably, *oda* transcript levels increased when putrescine was depleted, suggesting a negative feedback mechanism. These findings demonstrate that putrescine is not only an extracellular communal metabolite but is also vital for the cellular integrity and survival of *F. nucleatum* under anaerobic conditions.

**IMPORTANCE:** *Fusobacterium nucleatum* is a prominent member of the oral microbiota and has been linked to various human diseases, including periodontitis, preterm birth, and colorectal cancer. Despite its clinical significance, the metabolic requirements that support its growth and viability remain poorly understood. In this study, we identify the *oda* gene, which encodes ornithine decarboxylase, as essential for *F. nucleatum* survival due to its role in putrescine biosynthesis. We demonstrate that depletion of putrescine leads to severe growth and morphological defects, accompanied by widespread transcriptional changes. These findings reveal an underappreciated metabolic vulnerability and highlight the critical role of polyamine homeostasis in maintaining cellular integrity in this notorious anaerobe.

## INTRODCUTION

Polyamines are small, ubiquitous organic molecules containing two or more amine (–NH₂) groups that are maintained at micro-to millimolar concentrations across virtually all forms of life (1). In bacteria, the primary polyamines are putrescine and spermidine, which regulate diverse processes including gene expression, membrane integrity, motility, biofilm development, and intercellular as well as host–pathogen communication (2–5). Although polyamines are broadly required for growth, spermidine is indispensable in many species, with its depletion proving lethal (3). By contrast, putrescine is generally regarded as nonessential for viability, with *Ralstonia solanacearum*, a Gram-negative plant pathogen, being the only bacterium reported to strictly require putrescine for survival(3, 6).

Putrescine has also been detected in human gingival crevicular fluid (GCF), where it is thought to derive from subgingival dental plaque bacteria (7–9). Dental plaque harbors more than 400 microbial species, among which *Fusobacterium nucleatum* serves as a central organizer of community structure (10, 11). *F. nucleatum* is a Gram-negative, strictly anaerobic bacterium strongly associated with periodontitis and is distinguished by its ability to physically interact with a wide range of oral microbes—from early dental colonizers such as *Streptococcus gordonii* to late colonizers such as *Porphyromonas gingivalis* (12, 13). Beyond the oral cavity, *F. nucleatum* has been implicated in systemic diseases, including adverse pregnancy outcomes and colorectal cancer (14, 15). This species exhibits extensive genetic and phenotypic diversity and is currently divided into four subspecies (*nucleatum, vincentii, polymorphum,* and *animalis*), which have recently been proposed to represent distinct species(16–18).

Recent work demonstrated that *F. nucleatum* can import ornithine secreted by co-cultured *S. gordonii* and convert it into putrescine via ornithine decarboxylase (Oda) (19). The putrescine produced is then released extracellularly and has been shown to promote biofilm formation by *P. gingivalis*, suggesting that *F. nucleatum* contributes putrescine as a communal metabolite (19). Separately, *F. nucleatum* has been reported to secrete large amounts of putrescine during interactions with cancer cells, thereby promoting tumor progression (20). However, whether putrescine biosynthesis is required for the normal physiology and viability of *F. nucleatum* itself remains unknown.

In this study, we addressed this question by investigating the role of *oda* in *F. nucleatum*. We found that *oda* is essential, as deletion was only possible when a functional copy was supplied in trans. This essentiality was further confirmed using CRISPR interference and by generating a conditional mutant dependent on exogenous putrescine. Depletion of putrescine led to severe physiological defects, including filamentation, a conditional lysis phenotype, and widespread transcriptional dysregulation. Collectively, these findings demonstrate that *oda* and putrescine biosynthesis are indispensable for the viability of *F. nucleatum*.

## RESULTS

### Expression of *oda* peaks during mid-logarithmic growth

Defining a gene expression pattern during bacterial growth provides important insights into the gene’s function (21). The *oda* gene encodes ornithine decarboxylase/arginase, which catalyzes the conversion of ornithine to putrescine in *F. nucleatum* (22). Genomic analysis revealed that *oda* is organized as a monocistronic transcriptional unit (Fig. 1A). Although previous studies have shown that *oda* expression is markedly induced during co-culture with other oral bacteria (19, 23), its expression profile under monoculture conditions remains undefined.

**Figure 1.**
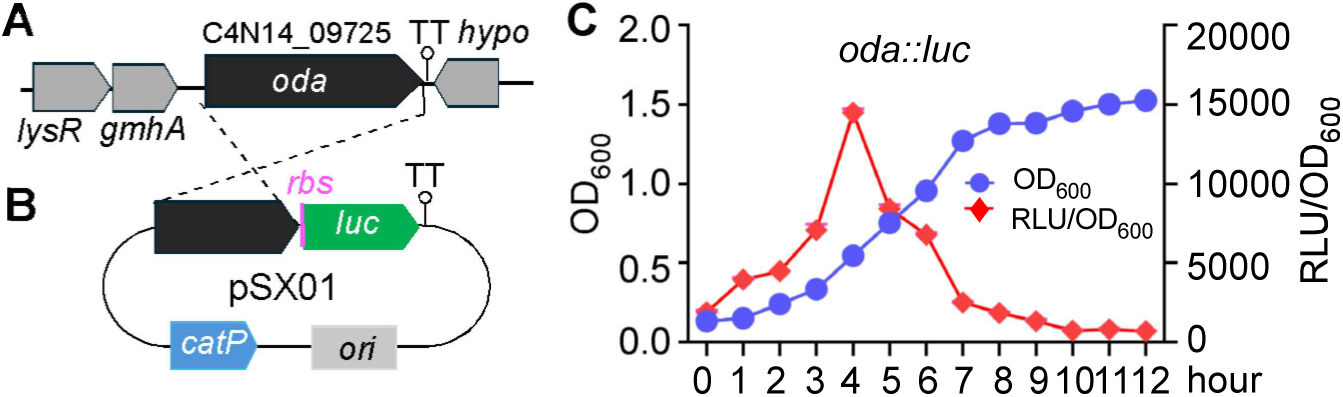
Expression of *oda* during growth of *Fusobacterium nucleatum*. **(A)** Genomic organization of the *oda* locus, showing it as a monocistronic transcriptional unit. **(B)** Construction of the *oda::luc* reporter plasmid (pSX01) and chromosomal integration into strain ATCC 23726. Features include TT (transcriptional terminator), *catP* (chloramphenicol/thiamphenicol resistance gene), and *ori* (origin of replication in *E. coli*). **(C)** Luciferase activity of the reporter strain during growth. The *oda* expression peaked at mid-log phase and declined in the stationary phase. Luminescence (RLU) was normalized to cell density (OD₆₀₀). Data represent means ± SD from three independent biological experiments.

To address this, we constructed a luciferase-based reporter plasmid, pSX01, in which the *luc* gene encoding Luciola red luciferase was fused to the 3′ end of *oda* (Fig. 1B). This construct was introduced and integrated into the chromosome of *F. nucleatum* strain ATCC 23726, generating a stable reporter strain. Luciferase activity was measured as a proxy for *oda* transcription across the growth curve. We observed a gradual increase in *oda* expression during early exponential growth, with luciferase activity peaking at mid-log phase before declining sharply (Fig.1C). By the stationary phase, *oda* expression was nearly undetectable (Fig. 1C). These results indicate that *oda* is most highly expressed during active cell proliferation, suggesting that putrescine biosynthesis is closely coupled to the growth phase and metabolic demands of *F. nucleatum*.

### *Oda* is essential in *F. nucleatum*

To investigate whether the *oda* gene function is related to the growth of *F. nucleatum*, we attempted to generate an in-frame deletion mutant using a two-step homologous recombination strategy previously developed in our lab (24). In the first step, two similarly sized DNA fragments flanking the *oda* coding region were cloned into the suicide vector pBCG02, which carries the HicA toxin gene as a counterselection marker (Fig. 2A) (24). The resulting plasmid, pBCG02-*oda*, was introduced into competent cells of strain ATCC 23726 and integrated into the chromosome through a single-crossover event at one of the homologous arms (Fig. 2B). Integrants were selected on thiamphenicol. These integrants represent merodiploid (*oda*/Δ*oda*) strains, in which a second homologous recombination event can occur spontaneously between the duplicated flanking regions. This event results in the excision of the plasmid and yields either a clean *oda* deletion mutant or a wild-type revertant, with equal probability (Fig. 2B). Since this second crossover occurs at a low frequency, we applied a toxic protein HicA-based counterselection to enrich for double-crossover recombinants by eliminating plasmid-retaining cells in the second step. Despite screening more than 60 colonies, we failed to recover any *oda* deletion mutants; all isolates retained the wild-type allele (Fig. 2C). These results strongly suggest that *oda* is essential for the viability of *F. nucleatum* under our tested growth conditions.

**Figure 2.**
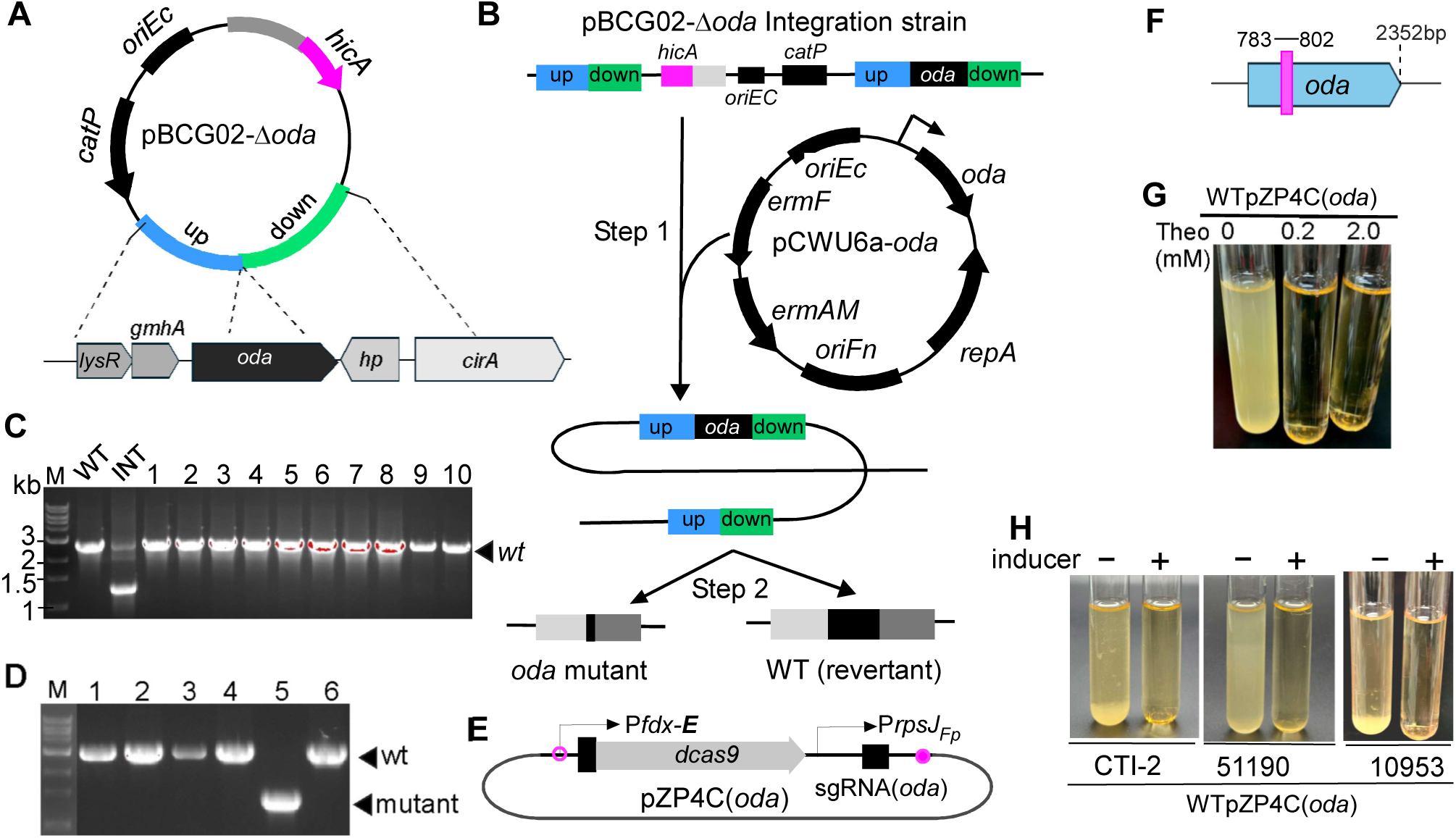
The *oda* gene is essential for *Fusobacterium nucleatum*. **(A)** Schematic of the pBCG02-Δ*oda* deletion construct designed for in-frame removal of the *oda* coding region. **(B)** Diagram of the allelic exchange strategy using pBCG02-Δ*oda*, which proceeds via a two-step recombination process in the presence of a second functional copy of *oda* carried on the expression plasmid pCWU6a-*oda*. **(C)** PCR screening of double-crossover candidates (>60 colonies examined; 10 representative colonies shown) revealed only the wild-type allele, indicating that *oda* could not be deleted. Int, integrated strain representing an *oda/Δoda* merodiploid. **(D)** In the presence of pCWU6a-*oda*, which provides an ectopic copy of *oda* under its native promoter, viable Δ*oda* mutants were successfully recovered following HicA-based counterselection, as confirmed by PCR. **(E)** Schematic of the CRISPRi system targeting *oda*, consisting of a theophylline-inducible dCas9 and a constitutively expressed sgRNA. **(F)** The specific 20-nt sequence within *oda* targeted by the sgRNA is shown (red line, position numbers indicated). **(G)** Growth of strain WTpZP4C(oda) was completely arrested upon theophylline induction (0.2–2.0 mM). (H) CRISPRi silencing of *oda* caused growth arrest across multiple *F. nucleatum* subspecies, including CTI-2 (*subsp. nucleatum*), ATCC 51190 (*subsp. vincentii*), and ATCC 10953 (*subsp. polymorphum*).

To confirm this essentiality, we tested whether deletion of the chromosomal *oda* allele could be achieved in a strain harboring an ectopic, functional copy of *oda*. We constructed a shuttle plasmid, pCWU6a-*oda*, in which *oda* is expressed under the control of its native promoter. Introduction of this plasmid into the pBCG02-*oda* integrant strain, followed by HicA-based counterselection, successfully yielded viable *oda* deletion mutants (Fig. 2D). This result confirms that the inability to delete *oda* in the wild-type background was due to its essentiality and that a functional copy provided in trans is sufficient to support cell viability.

To independently validate these findings, we utilized a riboswitch-regulated CRISPR interference (CRISPRi) system previously developed in our lab for studying essential genes in *F. nucleatum* (25). This system is based on a catalytically inactive *Streptococcus pyogenes* Cas9 protein (dCas9), which is specifically guided by a constantly expressed single-guide RNA (sgRNA) to the target gene (26). Upon induction by theophylline, dCas9 is expressed and binds to the sgRNA-specified site within the gene, blocking transcription by sterically hindering RNA polymerase progression. We identified a 20-nt target sequence within *oda* and constructed the corresponding CRISPRi plasmid, pZP4C(*oda*) (Fig. 2E–F). This plasmid was introduced into strain ATCC 23726 to generate the WTpZP4C(*oda*) strain. Upon induction with as little as 0.2 mM theophylline, the strain exhibited complete growth arrest (Fig. 2G), indicating effective transcriptional silencing and supporting the essentiality of *oda*.

To determine whether *oda* is essential across genetically diverse backgrounds, we introduced pZP4C(*oda*) into three additional *F. nucleatum* strains representing distinct subspecies: CTI-2 (subsp. *nucleatum*), ATCC 51190 (subsp. *vincentii*), and ATCC 10953 (subsp. *polymorphum*). In all three strains, theophylline-induced expression of dCas9 led to complete growth inhibition, mirroring the response observed in ATCC 23726. These results indicate that *oda* is universally essential across multiple *F. nucleatum* subspecies.

### Putrescine is essential for the growth of *F. nucleatum*

The *oda* gene encodes a bifunctional enzyme with an N-terminal ornithine decarboxylase (ODA) domain responsible for converting ornithine into putrescine, and a C-terminal arginase (ARG) domain (Fig. 3A, 3B). However, due to sequence divergence, the arginase domain is non-functional (27). Therefore, putrescine is presumed to be the primary product of *oda*-encoded activity.

**Figure 3.**
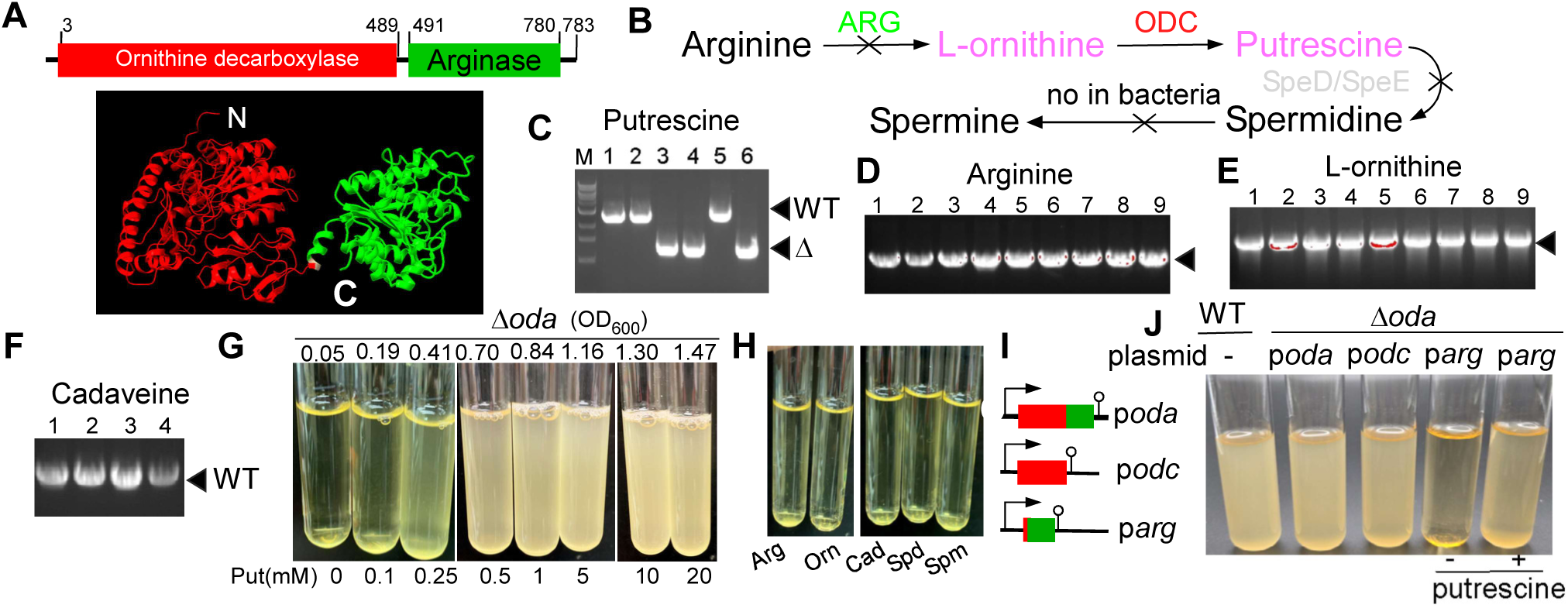
**Putrescine is essential for the growth of *Fusobacterium nucleatum***. **(A)** Domain organization of Oda, consisting of an N-terminal ornithine decarboxylase (ODC) domain and a nonfunctional C-terminal arginase-like (ARG) domain; a representative structural model was generated with AlphaFold. **(B)** Proposed polyamine biosynthetic pathways in *F. nucleatum*. Black X indicates missing enzymes that block the corresponding conversions. **(C–F)** Δ*oda* mutants were recovered only when plates were supplemented with putrescine (Put), as confirmed by PCR. Supplementation with arginine(Arg), ornithine(Orn), cadaverine(Cad), spermidine (Spd), or spermine(Spm) failed to support deletion, and only wild-type revertants were obtained. **(G–H)** Growth of the conditional Δ*oda* mutant was strictly dependent on exogenous putrescine in a dose-dependent manner, with no rescue by alternative metabolites. **(I)** Schematic of complementation plasmids: pBCG11-oda (full-length), pBCG11-odc (ODC domain), and pBCG11-arg (ARG domain). **(J)** Complementation with plasmids expressing full-length Oda or the ODC domain restored growth in the absence of putrescine, whereas expression of the ARG domain alone did not.

Notably, *F. nucleatum* lacks the canonical polyamine biosynthesis genes *speD* and *speE,* which are required for the synthesis of spermidine in *E. coli*, and, like most bacteria, cannot synthesize spermine *de novo* (2, 3). These facts support the hypothesis that *oda* essentiality may arise from a strict requirement for putrescine. If this hypothesis holds, supplying exogenous putrescine should allow isolation of *oda* deletion mutants from double-crossover recombinants following HicA-based counterselection.

Indeed, when screening plates were supplemented with 20 mM putrescine, 3 out of 6 randomly selected colonies were confirmed as *oda* deletion mutants (Fig. 3C). In contrast, supplementation with arginine, ornithine, or other polyamines such as cadaverine, spermidine, or spermine failed to yield any *oda* deletion mutants; all screened colonies retained the wild-type allele (Fig. 3D–F, data not shown). The resulting *oda* conditional mutant could not grow in the absence of exogenous putrescine, and its growth was strictly dependent on putrescine in a dose-dependent manner (Fig. 3G). Again, Arginine, ornithine, cadaverine, spermidine, and spermine were unable to rescue growth (Fig. 3H).

Given that the arginase-like domain is non-functional, the essential function of *oda* should reside solely within the N-terminal ODA domain. To test this, we constructed three expression plasmids encoding (i) the full-length *oda* gene, (ii) the N-terminal ODC domain alone, and (iii) the C-terminal ARG domain alone (Fig. 3I). Each plasmid was introduced into the *oda* deletion mutant. Strains expressing either full-length *oda* or only the ODC domain were able to grow in the absence of putrescine. In contrast, the strain expressing only the ARG domain failed to grow without supplementation (Fig. 3J). Together, these results demonstrate that putrescine is essential for *F. nucleatum* viability and that the essential activity of *oda* is encoded entirely within its N-terminal ornithine decarboxylase domain.

### Putrescine is essential for maintaining normal cell morphology and membrane integrity

To investigate why putrescine is essential for the growth of *F. nucleatum*, we examined the morphology of the *oda* mutant under varying putrescine concentrations using light microscopy.

Two methods were used for sample preparation. In the first (unwashed), 2–3 μL of culture was directly spotted onto a glass slide and examined under a microscope. In the second (washed) method, 1 mL of culture was centrifuged, washed twice with sterile water, and then resuspended in 300 μL of sterile water. Subsequently, 2–3 μL was used for microscopy.

In unwashed samples grown with 20 mM putrescine, the Δ*oda* mutant cells displayed a normal rod-shaped morphology similar to that of wild-type cells (Fig. 4A & 4B). No significant differences were observed between washed and unwashed wild-type cells (Fig. 4A & insert). However, washed *oda* mutant cells grown in 20 mM putrescine appeared morphologically altered compared to their unwashed counterparts (Fig. 4B vs 4C). Strikingly, under low putrescine conditions (0.25 mM), unwashed *oda* mutant cells were markedly elongated (Fig. 4D). Upon washing, these cells became highly adhesive and formed dense aggregates that were difficult to disperse (Fig. 4E). When gently agitated in a 24-well plate, the washed cells grown in 0.25 mM putrescine formed large clumps, while those grown in 20 mM putrescine formed only small particles. In contrast, washed wild-type cells showed no aggregation (Fig. 4H).

**Figure 4.**
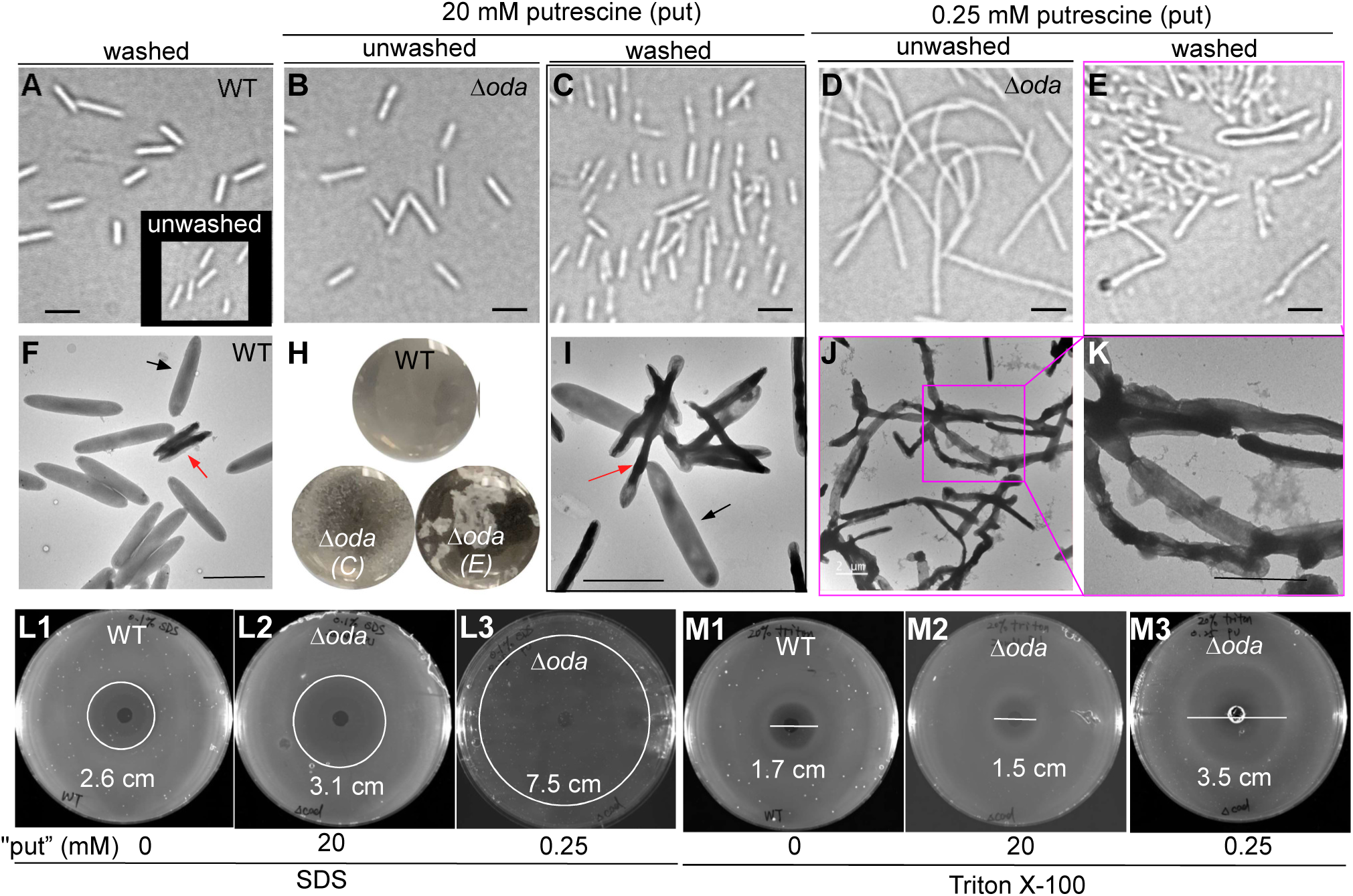
Putrescine is required for normal morphology and membrane integrity in *Fusobacterium nucleatum*. (A–B) Light microscopy of unwashed WT and Δ*oda* cells grown with 20 mM putrescine showed normal rod-shaped morphology. The inset in panel A shows washed WT cells. **(C)** Washed Δ*oda* cells grown with 20 mM putrescine displayed altered morphology compared to unwashed cells. **(D–E)** Under 0.25 mM putrescine, Δ*oda* cells became markedly elongated; washed cells formed dense aggregates that were difficult to disperse. Bars, 2 µm. **(F)** TEM of WT cells revealed mostly healthy cells with uniform staining (black arrow), along with a small fraction of cells showing signs of aging (red arrow). **(G–I)** Δ*oda* cells grown with 20 mM putrescine exhibited increased proportions of unhealthy cells, while those at 0.25 mM putrescine formed clumps and chains with widespread outer membrane rupture. **(J–K)** TEM images confirmed envelope disruption in Δ*oda* cells under low putrescine. Bars, 2 µm. **(L–M)** Detergent sensitivity assays. Both WT and Δ*oda* strains exhibited some susceptibility to SDS and Triton X-100; however, Δ*oda* cells grown with 0.25 mM putrescine were hypersensitive, displaying markedly larger inhibition zones compared to WT or Δ*oda* cells grown with 20 mM putrescine.

To further assess membrane structure, we performed negative staining with uranyl acetate and visualized the washed cells using transmission electron microscopy (TEM). Wild-type cells displayed two morphological populations: a predominant population (∼96%) of healthy cells with uniform gray staining and wider cell bodies, and a minor population (∼4%) of darker, narrower aging cells (Fig. 4F). In contrast, Δ*oda* cells grown in 20 mM putrescine showed ∼70% unhealthy cells, often attached to healthier neighbors (Fig. 4I). Cells grown in 0.25 mM putrescine displayed extensive clumping, often forming long chains, and exhibited widespread signs of outer membrane rupture (Fig. 4J & 4K).

These observations suggest that putrescine depletion disrupts cell division and impairs membrane integrity, contributing to abnormal morphology and cell lysis. To further test membrane vulnerability, we assessed detergent sensitivity using spot assays. Triton X-100, a nonionic detergent that disrupts the inner membrane, and SDS, an anionic detergent used to probe outer membrane barrier function in Gram-negative bacteria, were applied (28, 29). Compared to wild-type cells, *oda* mutant cells grown with 0.25 mM putrescine were highly sensitive to both detergents, displaying large zones of inhibition (Fig. 4L3 & 4M3). Cells grown with 20 mM putrescine showed only mild sensitivity to SDS and remained resistant to Triton X-100. (Fig. 4L1-2 & M1-2). Together, these findings suggest that the essentiality of putrescine in *F. nucleatum* may stem from its critical role in preserving normal cell morphology and membrane integrity.

### Limited putrescine supply remodels gene expression in *F. nucleatum*

To further investigate the essentiality of *oda*, we attempted to isolate spontaneous suppressor mutants that could compensate for the loss of *oda*. However, after plating Δ*oda* cells on rich medium without putrescine and incubating for 3–5 days, no colonies emerged, suggesting that putrescine’s role is indispensable and cannot be bypassed by simple secondary mutations.

To identify gene expression changes that might suggest a functional basis for putrescine essentiality, we performed transcriptomic analysis using RNA sequencing (RNA-seq). Total RNA was isolated from wild-type and Δ*oda* strains grown in the presence of either 20 mM or 0.25 mM exogenous putrescine. Differential expression analysis was performed using a two-fold cutoff (log₂ fold change ≥1 or ≤–1). When comparing Δ*oda* cells grown with 20 mM putrescine to wild-type, 186 genes were significantly upregulated and 174 were downregulated. Further comparison between Δ*oda* cells grown in 0.25 mM putrescine with WT revealed 331 upregulated and 353 downregulated genes (Fig. 5A & 5B and Tables S3 and S4). Cross-analysis of the two datasets identified 266 overlapping genes—108 consistently upregulated and 143 consistently downregulated—suggesting a core set of genes that respond in a dose-dependent manner to putrescine availability (Fig.5C, Fig. S1, and Tables S5, S6, S7).

**Figure 5.**
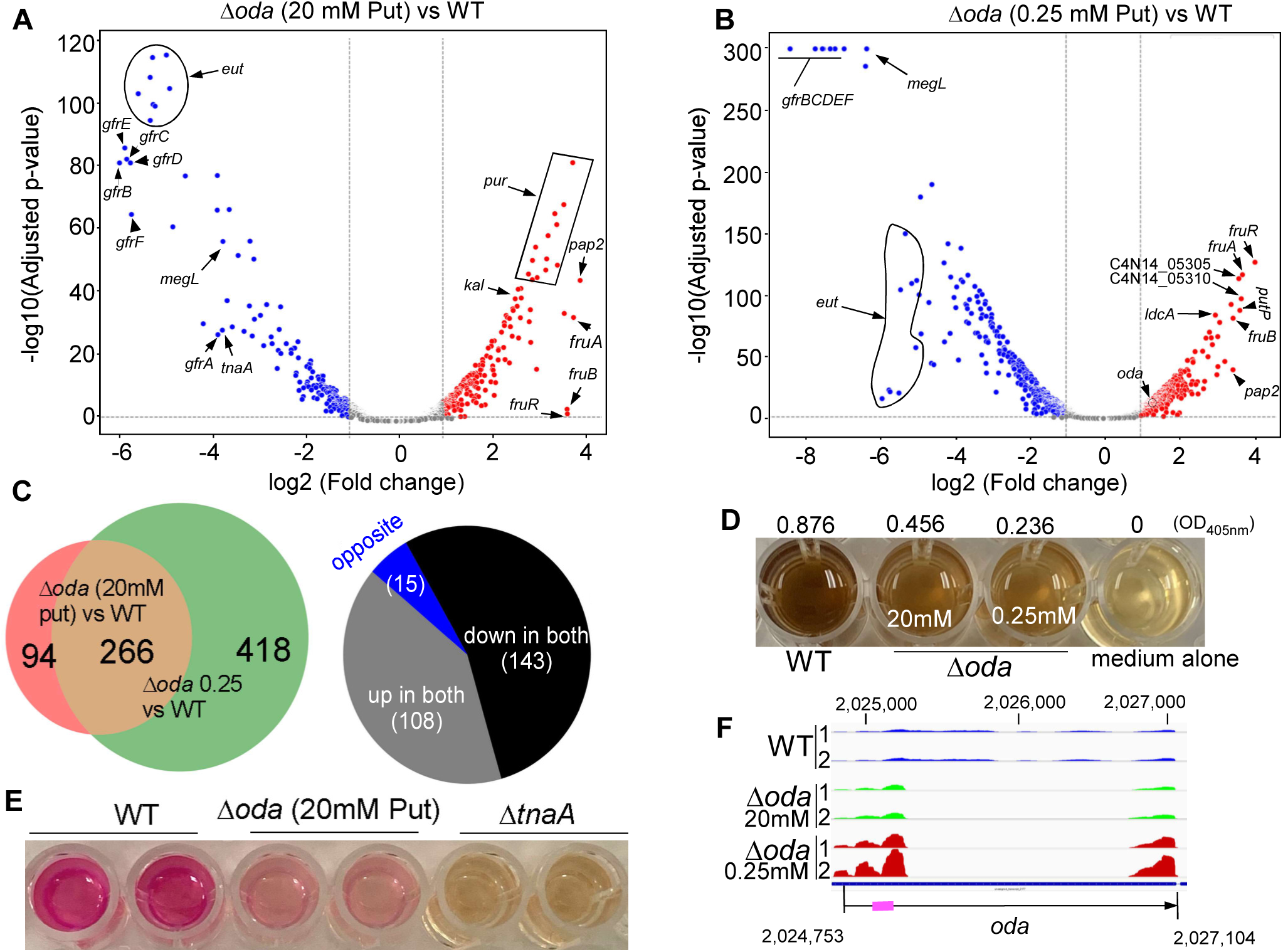
Transcriptomic remodeling of *Fusobacterium nucleatum* under putrescine limitation. (A–B) Volcano plots of differential gene expression determined by RNA-seq. Comparison of Δ*oda* (20 mM putrescine) versus WT identified up- and downregulated genes, while comparison of Δ*oda* cells grown with 0.25 mM versus 20 mM putrescine revealed an expanded set of transcriptional changes. **(C)** Venn diagram showing 266 overlapping genes consistently regulated under both conditions (108 upregulated, 143 downregulated, 15 oppositely regulated). **(D)** Downregulation of *megL*, encoding methionine γ-lyase, was functionally confirmed by reduced H₂S production, as measured by the bismuth sulfite assay. **(E)** Downregulation of *tnaA*, encoding tryptophanase, correlated with decreased indole production, verified using Kovac’s reagent assay. **(F)** RNA-seq coverage plots of the *oda* locus showed increased transcript signal in the undeleted 5′ and 3′ regions in Δ*oda* cells under low putrescine, suggesting feedback regulation of *oda* expression. The chromosomal location of *oda* in *F. nucleatum* ATCC 23726 is shown, with the region used for qRT-PCR highlighted by a thick red line.

Among the most strongly upregulated genes in Δ*oda* cells grown with 20 mM putrescine was *C4N14_07450*, encoding a PAP2-family phosphatidic acid phosphatase. These enzymes participate in lipid remodeling and membrane homeostasis by converting phosphatidic acid into diacylglycerol in bacteria (30). Expression of this gene increased even further under 0.25 mM putrescine (Fig. 5B), a result also confirmed by qRT-PCR (Fig. S1). Additional highly upregulated genes under putrescine-sufficient conditions included the *fruRBA* operon, which facilitates fructose uptake; multiple genes from the lysine degradation pathway; and a 10-gene operon spanning *C4N14_07225* to *C4N14_07270* involved in *de novo* purine biosynthesis. Interestingly, while the *pur* operon was induced at 20 mM putrescine, it was significantly downregulated when the putrescine concentration was reduced, suggesting a complex, concentration-dependent regulatory mechanism. One particularly responsive cluster, *C4N14_05285–C4N14_05320*, showed moderate induction at 20 mM putrescine and strong upregulation under 0.25 mM conditions. This operon includes *pelG*, which is involved in Pel polysaccharide biosynthesis and contributes to biofilm formation in several Gram-negative bacteria, including *Pseudomonas aeruginosa* (30, 31).

Conversely, the most downregulated genes included the *gfrABCDEF* operon, which encodes a mannose family phosphotransferase system-like involved in utilizing fructoselysine and glucoselysine (32). This operon was the most repressed in both conditions (Fig. 5A & 5B). Other significantly repressed targets included the *eut* operon (ethanolamine utilization) and *megL*, encoding methionine γ-lyase, a key enzyme in cysteine metabolism. Downregulation of *megL* was functionally confirmed using the bismuth sulfite assay, which revealed diminished H₂S production under low putrescine conditions (Fig. 5D & Fig.S1). We also observed downregulation of *tnaA*, which encodes tryptophanase and promotes the production of indole (33). Its expression was reduced in Δ*oda* cells grown with 20 mM putrescine compared to wild-type, as validated by the Kovacs’ reagent assay (Fig. 5E). However, further reduction of putrescine to 0.25 mM did not lead to additional downregulation. Interestingly, a distinct response was observed for the *C4N14_01755–C4N14_01760* operon, which was upregulated only at a concentration of 0.25 mM putrescine. This operon encodes *LdcA* (C4N14_01755), a muramoyltetrapeptide carboxypeptidase implicated in peptidoglycan recycling (34), and *PutP* (*C4N14_01760*), a sodium/proline symporter critical for L-proline uptake. These genes may reflect a stress-adaptive response to severe polyamine limitation.

Unexpectedly, *oda* expression itself was upregulated in Δ*oda* cells under 0.25 mM putrescine compared to 20 mM, despite the central region of the gene being deleted (∼1,650 bp) (Table S3 & S4). RNA-seq coverage plots revealed increased signal in the undeleted 5′ and 3′ regions, and qRT-PCR targeting the 5′ region confirmed this pattern (Fig. 5F and S1). These data suggest that *oda* expressions are negatively regulated by intracellular putrescine levels, possibly via feedback inhibition. Collectively, these findings demonstrate that putrescine limitation elicits a widespread transcriptional response in *F. nucleatum*, affecting genes involved in membrane remodeling, metabolic adaptation, biofilm development, and stress response. These regulatory shifts provide molecular insight into why putrescine biosynthesis is essential for viability and highlight its role in maintaining cellular homeostasis under nutrient-limited conditions.

## DISCUSSION

*F*. *nucleatum* is a keystone member of polymicrobial dental plaque, playing a central structural role in bridging early and late colonizers (11). This bridging capacity is attributed to its elongated morphology and robust coaggregation behavior, which enable it to mediate spatial organization and nutrient exchange within the biofilm community. One compelling example of metabolic cross-feeding is its ability to import ornithine secreted by neighboring bacteria and convert it into putrescine via the action of ornithine decarboxylase, encoded by the *oda* gene (19). The resulting putrescine is released extracellularly, where it promotes biofilm formation by species such as *P. gingivalis* . Furthermore, secreted putrescine has been implicated in tumor-promoting interactions between *F. nucleatum* and host cells (20). While these studies highlight the ecological significance of putrescine as a shared metabolite, its intrinsic role in the physiology of *F. nucleatum* has remained undefined. In this study, we demonstrate that putrescine is essential for the viability of *F. nucleatum* and that *oda*, the gene responsible for its biosynthesis, is indispensable. Genetic deletion of *oda* was only possible when complemented in trans, and its repression via CRISPR interference led to complete growth arrest. Putrescine depletion caused severe physiological consequences, including impaired cell division, filamentation, membrane destabilization, osmotic lysis, and widespread changes in gene expression.

Polyamines are widely conserved across bacteria, where they modulate diverse cellular functions, including gene regulation, membrane dynamics, and interspecies signaling (2–4) . While spermidine is essential in many organisms, putrescine has rarely been found to be indispensable, with *R. solanacearum* being the only previously reported case (6). Most bacteria synthesize both putrescine and spermidine, allowing one to compensate for the absence of the other. However, *F. nucleatum* lacks the canonical spermidine biosynthesis genes (*speD*, *speE*), suggesting that putrescine is the predominant (35), if not sole, polyamine in this organism. Its absence likely creates a unique metabolic bottleneck not easily circumvented by alternative pathways.

The essentiality of putrescine in *F. nucleatum* appears to be closely linked to its function in preserving cell morphology and envelope integrity. Under putrescine-limited conditions, cells of strain ATCC 23726—which normally exhibit a short, rod-shaped morphology—become markedly elongated and form chain-like structures, indicative of impaired cell division (Fig. 4A-D). These morphological abnormalities are accompanied by increased susceptibility to hypoosmotic stress, resulting in rapid cell lysis upon resuspension in water (Fig. 4). Transmission electron microscopy further revealed extensive membrane disruption and structural defects in putrescine-depleted cells (Fig. 4J & K). Consistent with compromised envelope integrity, these cells also displayed heightened sensitivity to the detergents Triton X-100 and SDS (Fig. 4L1-M3).

In certain anaerobic Gram-negative bacteria such as *Veillonella* spp. and *Selenomonas ruminantium*, polyamines like cadaverine and putrescine are covalently attached to the cell wall— linking D-glutamic acid residues in peptidoglycan stem to outer membrane proteins such as Mep45 (OmpM) (36–39).This linkage is important for normal growth, although it is not essential for viability(37). These bacteria lack the *lpp* gene, which encodes Braun’s murein lipoprotein, a structural anchor in many Gram-negative species (40). Interestingly, *F. nucleatum* also lacks *lpp* and contains D-glutamic acid in its peptidoglycan stem (L-Ala-D-Glu-L-Lan-D-Ala) (40, 41), and expresses multiple OmpM homologs, including FomA, which appears to be essential for cell viability as well (Wu et al., unpublished data). These parallels suggest that *F. nucleatum* may likewise incorporate putrescine into its envelope, potentially bridging the peptidoglycan and outer membrane to reinforce structural stability. Supporting this hypothesis, transcriptomic data revealed a strong induction of *C4N14_07450* (a PAP2-family phosphatase) and *C4N14_01755* (LdcA), both of which are associated with cell wall metabolism, under putrescine-limited conditions. The Δ*oda* mutant also showed increased susceptibility to osmotic stress, lysing rapidly upon washing and resuspension in water. This fragility was accompanied by upregulation of a betaine uptake operon (*C4N14_02945–C4N14_02950*) (Table S3 & S4), suggesting that *F. nucleatum* mounts an osmoprotective response in the face of polyamine depletion. Betaine, a known compatible solute, may partially buffer the cytoplasm against osmotic shock, but this response alone is insufficient to restore cell viability.

Although supplementation with 20 mM putrescine restored growth and morphology in Δ*oda* cells, it did not fully replicate the wild-type phenotype (Fig. 4). Several factors may underlie this incomplete rescue. First, intracellular putrescine concentrations may remain suboptimal; in *E. coli,* cellular levels range from 10 to 30 mM (41). Second, *F. nucleatum* may lack efficient high-affinity transport systems for polyamines. Although it encodes a putative *potABCD* transporter, these genes (*C4N14_01555–01565* and *C4N14_09145*) were not induced in the Δ*oda* mutant, suggesting limited uptake capacity under the tested conditions. Third, Oda may catalyze the formation of additional, yet unidentified metabolites from ornithine or alternative substrates that are critical for cellular homeostasis. Lastly, putrescine may require post-synthetic modifications to perform its essential functions. The conserved but enzymatically inactive arginase-like domain in Oda, present across all sequenced *F. nucleatum* genomes, raises the possibility of a non-catalytic structural role—perhaps in modifying putrescine within the cell cytoplasm. Ongoing studies in our lab are exploring these possibilities.

Transcriptomic analysis revealed that putrescine limitation results in a global remodeling of gene expression. A core set of 266 genes showed dose-dependent regulation in Δ*oda* cells (Table S5& S6). Upregulated pathways included lipid metabolism (PAP2), fructose uptake (*fruRBA*), lysine degradation, and purine biosynthesis. Interestingly, co-upregulation of *fruRBA* and the lysine degradation operon under low putrescine levels contradicts their previously observed inverse regulation, suggesting a unique transcriptional landscape under polyamine stress. Downregulated genes included *gfrABCDEF* (carbohydrate transport), the *eut* operon (ethanolamine utilization), *tnaA* (indole production), and *megL* (hydrogen sulfide biosynthesis), highlighting shifts in carbon, nitrogen, and sulfur metabolism. Notably, a specific operon encoding LdcA (a peptidoglycan hydrolase) and PutP (a proline transporter) was strongly induced only under severe putrescine limitation, implying a stress adaptation mechanism tied to envelope maintenance and amino acid acquisition.

Surprisingly, *oda* expression itself was elevated in Δ*oda* cells under low putrescine, despite the central deletion. Increased transcript levels in the 5′ and 3′ regions of *oda* suggest the presence of a negative feedback loop wherein intracellular putrescine represses its own biosynthesis. No regulatory protein controlling *oda* has been reported in bacteria. We hypothesize the existence of a yet-unidentified transcriptional regulator that responds to intracellular putrescine levels by modulating *oda* transcription in *F. nucleatum*. At low putrescine concentrations, this regulator may activate expression; upon sufficient accumulation, putrescine binding could suppress its activity, leading to autoregulation. This model is consistent with the observed growth-phase-dependent expression of *oda*, which peaks during exponential growth and diminishes in the stationary phase. In summary, our findings establish *oda* and putrescine biosynthesis as essential for *F. nucleatum* survival. These results uncover a previously underappreciated role for putrescine in maintaining membrane architecture, osmotic resilience, and global gene expression in this organism. Beyond its well-recognized ecological and pathogenic functions, putrescine is vital for *F. nucleatum*’s own cellular integrity. This work deepens our understanding of polyamine biology in anaerobic bacteria, providing new insights into targeting polyamine metabolism in oral biofilms and cancer-associated microbiomes.

## MATERIALS AND METHODS

### Bacterial strains and growth conditions

The bacterial strains and plasmids used in this study are listed in **Table S1**. *Fusobacterium nucleatum* strains were cultured in tryptic soy broth (TSB) supplemented with 1% Bacto Peptone and 1 mM freshly prepared cysteine (referred to as TSPC), or on TSPC agar plates. Anaerobic cultivation was performed in a Coy anaerobic chamber (Coy Laboratory Products) filled with a gas mixture consisting of N₂, H₂, and CO₂. Hydrogen levels were continuously monitored and maintained above 2% using a portable H2 sensor placed inside the chamber. *Escherichia coli* strains were grown aerobically in Luria–Bertani (LB) broth or on LB agar plates. When required, antibiotics were added at the following final concentrations: chloramphenicol (15 μg/mL), thiamphenicol (5 μg/mL), erythromycin (300 μg/mL for LB agar; 50 μg/mL for LB broth), and clindamycin (1 μg/mL)

### Plasmids construction

All primers used in this study were custom-synthesized by Sigma-Aldrich and are listed in Table S2. New plasmids were constructed using Gibson assembly cloning, following the manufacturer’s protocol with the NEBuilder HiFi DNA Assembly Master Mix (New England Biolabs, E2621L). The integrity of all constructs was verified by DNA sequencing.

i. **pCM-galK-*oda::luc*** — This plasmid was designed to transcriptionally fuse the *luciola* red luciferase gene (*luc*) to the 3′ end of *oda*. A 2,629-bp DNA fragment, encompassing part of the *oda* promoter and the entire coding sequence (from the start codon to the stop codon), was amplified using primers oda-F and oda-R. The *luc* gene was PCR-amplified from plasmid pBCG06 (42) using primers luc-F and luc-R. The two fragments were fused via overlap extension PCR to generate the *oda::luc* fusion. This product was then inserted into the suicide plasmid pCM-galK, which had been linearized by PCR using primers 2pCM-galK-F and 2pCM-galK-R. Gibson assembly was used to complete the construction.
ii. **pBCG02-Δ*oda*** — The *oda* gene encodes a 783–amino acid protein. We aimed to design a construct to delete the region corresponding to amino acids 174–654. To achieve this, ∼1.2-kb DNA fragments upstream and downstream of *oda* were PCR-amplified using primer pairs oda-up-F/oda-up-R and oda-dn-F/oda-dn-R, respectively. The two flanking fragments were assembled into the pBCG02 backbone (33), which had been PCR-linearized with primers BCG02Mch-F and BCG02Mch-R, using Gibson assembly.
iii. **pCWU6a-*oda*** — The shuttle plasmid pCWU6 contains a unique EcoRV restriction site within the middle of the *catP* gene. To construct pCWU6a-oda, pCWU6 was linearized by EcoRV digestion. An erythromycin resistance cassette (comprising *ermF* and *ermAM*) was PCR-amplified from plasmid pG106 (43) using primer pair 6a-ermF/6a-ermR. The resulting amplicon was inserted into the linearized pCWU6 via Gibson assembly. Recombinant plasmids were selected on LB agar plates supplemented with 300 μg/mL erythromycin. To construct pCWU6a-oda, pCWU6a was further linearized by PCR using primers pCWU6a-F/pCWU6a-R. The *oda* gene along with its native promoter was amplified from genomic DNA using primers oda-G-F/oda-G-R, and the resulting amplicon was cloned into the linearized pCWU6a backbone via Gibson assembly, yielding the final construct pCWU6a-oda.
iv. **pZP4C(*oda*)** — A 20-nt target sequence for the *oda* gene (5′-GATATGATGGAGAAGTTGTA-3′) was selected for CRISPR interference. This sequence was incorporated into the forward primer to create P1(*oda*), which was used together with primer P2 to amplify the sgRNA scaffold region using pZP4C (25) as the DNA template. The resulting PCR product was cloned into MscI/NotI-digested pZP4C-mCherry via Gibson assembly.
v. **p*oda*, p*odc* and p*arg*** — The *oda* gene together with its native promoter was PCR-amplified using primer pair com-odaF/com-odaR, and the resulting amplicon was cloned into pBCG11, a small shuttle plasmid. For cloning, pBCG11 was linearized by PCR with primers pBCG11-F/pBCG11-R, and the linearized vector was assembled with the amplicon by Gibson assembly to generate p*oda*. To express only the Odc domain of Oda, p*oda* was subjected to inverse PCR using primers oda1-502-F/oda1-502-R to delete the region encoding amino acids 503–784. The inverse PCR product was phosphorylated with T4 polynucleotide kinase and ligated using Quick DNA ligase (44), yielding p*odc*. Following a similar strategy, p*arg* was generated to express a truncated variant of Oda containing amino acids 1–20 and 490–783. For this construct, inverse PCR was performed using primers oda(arg)-F/oda(arg)-R.

### Initial attempt to make an in-frame deletion of *oda*

A published procedure employing *hicA* as a counterselection marker was adapted to generate a non-polar, in-frame deletion of *oda* in *F. nucleatum*. Briefly, the deletion plasmid pBCG02-Δ*oda* was introduced into competent cells of strain ATCC 23726 by electroporation. Integration of the plasmid into the chromosome via homologous recombination was selected on TSPC agar plates containing 5 µg/mL thiamphenicol. To screen the second recombination event, which can either excise the vector with restoration of the wild-type allele or result in deletion of the target gene, transformants were plated on medium supplemented with 2 mM theophylline, the inducer of HicA toxicity. PCR screened colonies exhibiting the expected phenotype (toxin-resistant and thiamphenicol-sensitive) for the intended deletion. Approximately 80 colonies were examined; however, all retained the wild-type allele, and no in-frame deletion mutants were recovered. These results indicate that *oda* is essential under the tested conditions.

### Creation of a conditional *oda* deletion mutant

For the creation of a conditional mutant, the integrant strain carrying pBCG02-Δ*oda* was complemented in trans with pCWU6a-*oda*, which expresses *oda* from its native promoter. Transformants were selected on TSPC agar containing 5 µg/mL thiamphenicol and 1 µg/mL clindamycin. A single colony was inoculated into TSPC broth with clindamycin and grown overnight. The culture was diluted 1:100, and 100-µL aliquots were spread on TSPC agar plates supplemented with 2 mM theophylline and clindamycin. After three days of anaerobic incubation at 37°C, colonies were screened for loss of thiamphenicol resistance, and thiamphenicol-sensitive isolates were examined by PCR to assess deletion of the chromosomal *oda* locus.

To test whether exogenous metabolites could compensate for the loss of *oda*, the integrant strain was cultured overnight in TSPC broth supplemented with 20 mM of arginine, ornithine, or individual polyamines in the absence of antibiotics. The cultures were then diluted at 1:100, and 100-µL aliquots were plated on TSPC agar containing 2 mM theophylline and 20 mM of the corresponding supplement. After three days of anaerobic incubation at 37 °C, colonies were screened for thiamphenicol sensitivity and PCR-verified putative deletion mutants.

### Luciferase assays

The reporter strain WTpCM-galK-*oda::luc* was cultured overnight in TSPC broth supplemented with 5 µg/mL thiamphenicol. Cells were harvested, washed twice with fresh TSPC, and used to inoculate fresh medium at a 1:10 dilution. Cultures were incubated anaerobically, and samples were collected at hourly intervals over a 12-h period. At each time point, three 150-µL aliquots were transferred into a 96-well microplate (#3917, Corning®) and mixed with 25 µL of 1 mM D-luciferin (Molecular Probes) prepared in 100 mM citrate buffer (pH 6.0). The mixtures were pipetted vigorously 15 times under aerobic conditions to ensure oxygen exposure and substrate mixing. Luminescence was measured using a GloMax Navigator Microplate Luminometer, and culture growth was monitored in parallel by measuring the optical density at 600 nm (OD₆₀₀). All assays were performed in triplicate and repeated independently on three separate occasions.

### CRISPRi assay

Fusobacterial strains carrying the CRISPRi plasmid were grown overnight in TSPC medium supplemented with 5 µg/mL thiamphenicol without inducer. Overnight cultures were then diluted 1:1000 into fresh TSPC medium containing the indicated concentrations of theophylline and incubated for 22 h.

### Autoaggregation assay and Electron microscopy

Wild-type (WT) and Δoda strains of *F. nucleatum* were cultured anaerobically for 14 hours in TSPC medium supplemented with either 20 mM or 0.25 mM putrescine. After incubation, the optical density at 600 nm (OD₆₀₀) was measured. Equivalent amounts of cells from each culture (normalized by OD₆₀₀) were harvested by centrifugation, washed twice with sterile distilled water, and resuspended in an equal volume of water. A 0.4-mL aliquot of each cell suspension was transferred into individual wells of a 24-well tissue culture plate. The plates were gently agitated by hand on the benchtop for 1 minute to initiate autoaggregation. Images were recorded immediately thereafter.

For electron microscopy, 10 µL of each bacterial suspension was applied onto carbon-coated nickel grids and allowed to settle. Grids were negatively stained with 0.1% (w/v) uranyl acetate and briefly rinsed with sterile water. Samples were visualized using a JEOL JEM-1400 transmission electron microscope.

### SDS and Triton X-100 sensitivity assay

Sensitivity to membrane-disrupting agents was assessed using an agar diffusion assay. Wells of 0.3 mm diameter were punched into TSPC agar plates using a sterile pipette tip. Each well was filled with 50 µL of either 0.1% (w/v) SDS or 20% (v/v) Triton X-100. After allowing the detergents to diffuse into the agar, a 10-mL overlay of 1.2% TSPC soft agar was prepared. This overlay was mixed with a 100-µL aliquot of bacterial suspension from 14-hour cultures of wild-type and Δ*oda* strains grown in TSPC medium containing either 20 mM or 0.25 mM putrescine. Prior to mixing, OD₆₀₀ values were measured to normalize cultures, and equivalent cell numbers were used for all conditions. The mixture was poured over TSPC agar base plates that either contained or lacked 20 mM or 0.25 mM putrescine, matching the respective growth condition. Plates were incubated anaerobically at 37°C for 72 hours, after which zones of inhibition surrounding each well were recorded.

### Gene expression profiling by RNA-seq

An overnight culture of the Δ*oda* mutant was grown anaerobically in TSPC medium supplemented with 20 mM putrescine. The following day, cells were harvested by centrifugation, washed twice with fresh TSPC medium to remove residual putrescine, and resuspended in fresh TSPC at an OD₆₀₀ of 1.0. Cultures were then subcultured at a 1:20 dilution into TSPC medium containing either 20 mM or 0.25 mM putrescine and incubated anaerobically. After 10 hours of growth, cells were collected for RNA extraction. A 10-hour wild-type culture grown without putrescine was also prepared for comparison. RNA samples were submitted to Cancer Genomics Center at The University of Texas Health Science Center at Houston (CPRITRP240610). Total RNA was quality-checked using Agilent RNA 6000 Pico kit (#5067-1513) by Agilent Bioanalyzer 2100 (Agilent Technologies, Santa Clara, USA). RNA with an integrity number of greater than 7 was used for library preparation. Libraries were prepared from the rRNA depleted RNAs withNEBNext Ultra II Directional RNA Library Prep Kit for ILLumina (E7760L, New England Biolabs) and NEBNext Multiplex Oligos for illumina (E6609S, New England Biolabs) following the manufacturer’s instructions. The quality of the final libraries was examined using Agilent High Sensitive DNA Kit (#5067-4626) by Agilent Bioanalyzer 2100 (Agilent Technologies, Santa Clara, USA), and the library concentrations were determined by qPCR using Collibri Library Quantification kit (#A38524500, Thermo Fisher Scientific). The libraries were pooled evenly and subjected to paired-end 150-cycle sequencing on an Illumina NovaSeq System (Illumina, Inc., USA). Paired-end reads were processed, mapped, and analyzed for differential gene expression using DESeq2(45). For further downstream analysis and visualization, R/Bioconductor packages were used. A threshold of |log₂(fold change)| ≥ 1.0 was used to define differentially expressed genes under the tested conditions.

### Indole assay

Overnight cultures of WT and Δ*oda* (20 mM putrescine) were grown to the stationary phase (OD₆₀₀ ≈ 1.4). Supernatants were collected by centrifugation, and 100-µL aliquots were transferred into wells of a 96-well plate. An equal volume of Kovac’s reagent was added without mixing. Indole production was indicated by a pink-to-red color within 30 s, with intensity reflecting indole concentration. A *tnaA* mutant served as a negative control and remained yellow. The assay was performed independently at least three times.

### Detection of H**₂**S

The Δ*oda* mutant was cultured in TSPC medium supplemented with either 20 mM or 0.25 mM putrescine for 10 h, while the WT strain was grown in unsupplemented TSPC. Equivalent amounts of cells from each culture were harvested, resuspended in 400 µL TSPC without putrescine, and transferred into wells of a 24-well plate. Each suspension was mixed with an equal volume of bismuth buffer [0.4 M triethanolamine-HCl (pH 8.0), 10 mM bismuth(III) chloride, 20 µM pyridoxal 5-phosphate monohydrate, 20 mM EDTA, and 40 mM L-cysteine] (46). After incubation at 37°C for 30 min, H₂S production was quantified by measuring absorbance at 405 nm. Results represent three independent experiments performed in duplicate.

## Data Availability

The RNA-seq raw data were deposited in the NCBI Sequence Read Archive (SRA) database with the accession number of PRJNA1311141.

## AUTHOR CONTRIBUTIONS

C.W. and S.X. conceived and designed all experiments. S.X., B.G., A.F., and C.W. performed all experiments. S.X., B.G., and C.W. analyzed data. C. W. S. X. wrote the manuscript with contributions and approval from all authors.

## ACKNOWLEDGMENTS

This work was supported by NIDCR grants DE030895 and DE034542 awarded to C.W. We thank the technical support from the Cancer Prevention and Research Institute of Texas (CPRITPR240610).

